# Spatiotemporal Atlas of Pro-Inflammatory (NF-κB) and Anti-Inflammatory (STAT6) Signalling Using Reporter Mice during mRNA Vaccination

**DOI:** 10.64898/2026.01.29.702227

**Authors:** Electra Brunialti, Alessia Panzeri, Nicoletta Rizzi, Alessandro Villa, Clara Meda, Chiara Sfogliarini, Stefano Persano, Federico Arlati, Maria L Guevara Lopez, Cristina Martelli, Gaetano Cannavale, Monica Rebecchi, Clara Di Vito, Letizia Venturini, Luisa Ottobrini, Domenico Mavilio, Susanna Fagerholm, Saverio Minucci, Elisabetta Vegeto, Paolo Ciana

## Abstract

The immunization process unfolds through a precisely orchestrated sequence of innate and adaptive events across distinct anatomical sites. Although many mechanisms underlying vaccination are well described, most vaccines have been developed empirically, partly due to the lack of tools enabling rapid, organ-specific analysis of immune activation. To address this gap, we developed and validated a novel STAT6 reporter mouse enabling dynamic *in vivo* whole-body imaging and *ex vivo* analysis of STAT6-mediated anti-inflammatory signalling, and combined it with an established NF-κB reporter model to dissect immune activation induced by two LNP-encapsulated mRNA vaccines encoding the same antigen but differing in RNA chemistry (unmodified versus N¹-methyl-pseudouridine (m¹Ψ)–modified).

This dual-reporter system enabled the creation of a spatiotemporal atlas of vaccine-induced signalling, revealing chemistry-dependent immune dynamics and identifying the liver as a predominant early hub for both NF-κB and STAT6 activity following systemic administration. Integration with antibody measurements demonstrated that early STAT6 activation followed by rapid signal resolution—rather than prolonged NF-κB–mediated inflammation—correlated with robust humoral responses, suggesting that monitoring NF-κB and STAT6 dynamics could provide predictive insight into vaccine immunogenicity.

Together, these findings establish NF-κB and STAT6 reporter mice as rapid *in vivo* screening tools for the early assessment of vaccine immunogenicity and performance. By enabling dynamic, organ-resolved immune profiling, this approach paves the way for more rational, mechanism-driven design of mRNA vaccines and underscores the importance of further investigating the effects of vaccines on the liver, both as a primary LNP target and as an immunologically tolerogenic organ.

## Introduction

The immunization process unfolds through a highly coordinated sequence of molecular and cellular events that progressively recruit innate and adaptive immune components across distinct anatomical niches ^1^. Immune activation against antigens begins with rapid innate sensing followed by the staged engagement of adaptive mechanisms. Within minutes, antigens are detected by pattern-recognition receptors (PRRs) expressed on innate immune sentinels. This immediate detection triggers inflammatory signalling cascades, cytokine release, and recruitment of additional effector cells, creating the microenvironment required for subsequent adaptive priming. Over the next 6–24 hours, antigen-presenting dendritic cells (DCs) undergo maturation and migrate to draining lymph nodes, where they present peptide–MHC complexes to naïve CD4⁺ and CD8⁺ T cells while delivering the co-stimulatory signals required for activation. Within 1–3 days, T cells bearing cognate T cell receptors undergo metabolic and transcriptional reprogramming that initiates rapid clonal expansion ^2–4^. Activated CD4⁺ T cells provide cytokines such as interleukin (IL)-2 that amplify CD8⁺ T-cell expansion, cytotoxic differentiation, and trafficking toward sites of antigen presence. In parallel, activated B cells interact with CD4⁺ T cells, receiving instructive signals mediated by CD40L, IL-4, and IL-21, which drive class switching and commitment either to early plasmablast differentiation and entry into germinal centers ^2–4^. The primary antibody response becomes detectable within 4–7 days, initially dominated by IgM production, followed by class-switch recombination and affinity maturation. Within germinal centres, B-cell clones undergo affinity-based selection, ultimately giving rise to long-lived plasma cells and memory B cells that sustain durable, high-affinity antibody production. After the primary response contracts over 2–4 weeks, antigen-specific memory T and B cells persist, providing a rapid and long-lasting response upon re-exposure ^5,6^.

Accordingly, successful immunization depends on the coordinated and temporally ordered engagement of molecular and cellular programs, governed by transcriptional regulators and finely tuned intercellular signalling networks involving cytokines, chemokines, and co-stimulatory molecules. The spatial and temporal control of these pathways across distinct anatomical compartments is essential for generating robust and long-lasting protective immunity while limiting pathological inflammation and preserving immune homeostasis^4^. Within this regulatory landscape, NF-κB and STAT (Signal Transducer and Activator of Transcription) signalling pathways function as key hubs integrating inflammatory cues with lymphocyte activation, proliferation, and differentiation^7^ ^8^.

NF-κB, a key mediator of pro-inflammatory responses, acts as an immediate-early transcription factor. It is activated within minutes of antigen sensing in peripheral tissues via PRR-dependent IκB degradation and nuclear translocation ^7^. This early activity drives the production of inflammatory cytokines, chemokines, and co-stimulatory molecules that coordinate DC maturation ^9,10^. In draining lymph nodes, sustained yet tightly regulated NF-κB activity over 6–24 hours further fine-tunes T-cell activation, thereby ensuring an effective adaptive immune response ^11,12^.

In parallel, STAT proteins direct the polarization of distinct T helper (Th) cell subsets. Among them, STAT6 mediates over 80% of IL-4-regulated genes, the key anti-inflammatory cytokine, and acts as a major driver of the transcriptional program defining Th2 differentiation^13^. STAT6 activation marks a later, instructive phase of adaptive immunity, driving IgE class switching and broader Th2-specific transcriptional responses^14^, while STAT6 signalling within B cells is essential for proper germinal center formation^8^. Importantly, dysregulation of NF-κB or STAT6 compromises both the magnitude and quality of immune responses, leading to insufficient T- and B-cell activation, impaired antibody production, and altered Th2 polarization, ultimately reducing vaccine efficacy or increasing the risk of excessive inflammation ^9,15,16^.

Despite extensive knowledge of immunological mechanisms, most vaccines have been developed empirically^4^, and the spatiotemporal dynamics of NF-κB and STAT6 *in vivo* remain poorly defined. Clarifying when and where these pathways are engaged could guide the rational design of vaccines and adjuvants to optimize immunity while limiting harmful inflammation.

In this context, the use of *in vivo* murine reporter systems represents a powerful experimental approach, as it allows for the dynamic and non-invasive analysis of transcription factor activation in living organisms ^17^. Leveraging this strategy, our lab has investigated the temporal and organ-specific functions of various transcription factors in both physiological and pathological contexts ^18–20^, including studies on pro-inflammatory responses using the NF-kB reporter mouse ^21^. However, the development of reporter systems capable of tracking complementary anti-inflammatory pathways would be essential to obtain a comprehensive view of immune regulatory processes.

To address these gaps, we first generated a novel reporter mouse model to investigate anti-inflammatory signalling mediated by STAT6 by optical *in vivo* imaging. We then developed *in vivo* imaging strategies that enable real-time, longitudinal monitoring of both NF-κB and STAT6 activation following immunization: we compared two mRNA vaccine constructs encoding the same antigen but differing in their chemical composition: an unmodified mRNA expected to strongly engage innate RNA sensors, and an N¹-methyl-pseudouridine (m¹Ψ)-modified mRNA designed to attenuate innate immune recognition while enhancing mRNA stability and translational efficiency^22–24^. By administering the vaccines intravenously to achieve systemic biodistribution, we generated a spatially and temporally resolved atlas of NF-κB and STAT6 activation across the body, providing a comprehensive framework to elucidate how vaccine chemistry influences immune signalling dynamics *in vivo*.

## Results

### Generation of a STAT6 reporter mouse line

To enable real-time monitoring of the anti-inflammatory pathway mediated by STAT6, we developed a dual-reporter system designed to track STAT6 transcriptional activity both *in vivo* and *ex vivo*. The reporter construct was based on a bicistronic design comprising a firefly luciferase gene (*luc2*) for *in vivo* bioluminescence imaging and a green fluorescent protein (GFP) gene for single-cell analysis, separated by an internal ribosome entry site (IRES) ^21^.

Upstream of the bicistronic cassette, we cloned a synthetic promoter containing two tandem copies of the validated STAT6 response element from the *p2xSTAT6-Luc2P* plasmid, generating the *p2xSTAT6-GFP-luc2* construct. The ability of this promoter configuration to report STAT6 activation was tested by transient transfection in RAW 264.7 macrophages. Cells were stimulated with cytokines known to activate STAT6-dependent transcription (IL-4 or IL-13, 20 ng/ml) ^25^, the anti-inflammatory cytokine IL-10 (20 ng/ml, acting through STAT3) ^26^, or the pro-inflammatory stimulus lipopolysaccharide (LPS, 2 µg/ml) (Figure 1A). Bioluminescence assays performed 24 hours after cytokine exposure, or 6 hours after LPS treatment, revealed a robust induction of luciferase activity following IL-4 and IL-13 stimulation, whereas IL-10 and LPS failed to elicit reporter activation. These results confirmed the specificity of the construct for STAT6-mediated signalling. The functionality of the GFP reporter was evaluated in HT-29 cells transiently transfected with the STAT6 construct together with a plasmid constitutively expressing tdTomato, used as a positive control for transfection efficiency. A clear GFP fluorescence was detected in tdTomato-positive cells following 24 h treatment with 20 ng/ml IL-4, whereas no significant GFP signal was observed in any cells treated with 100 ng/ml LPS or vehicle (water) (Figure 1B). These results demonstrate the proper functionality of both reporters. Encouraged by these results, we next generated the STAT6 reporter mice to investigate the modulation of anti-inflammatory signalling *in vivo*. Following a previously validated knock-in strategy routinely used in our laboratory, the reporter cassette was inserted into a defined transcriptionally active locus on mouse chromosome 7, which we have shown to support stable, position-independent expression ^21^. To insulate the transgene from chromatin position effects, the construct was flanked by matrix attachment region (MAR) sequences derived from the chicken lysozyme gene ^27^ (Figure 1C). In addition, the reporter cassette was preceded by a floxed STOP sequence (RNA polymerase II termination signal), allowing tissue-specific activation upon Cre-mediated recombination. Embryonic stem cells harbouring the correctly integrated construct were injected into blastocysts to generate chimeric mice carrying the floxed transgene, hereafter referred to as STAT6-STOP (Figure 1C). As expected, STAT6-STOP mice exhibited very low basal luciferase activity in *in vivo* imaging, confirming effective transcriptional blockade by the STOP cassette (Figure 1C and S1A). To obtain a constitutively active reporter line, STAT6-STOP mice were crossed with *B6.C-Tg(CMV-cre)1Cgn/J* mice, in which Cre recombinase is expressed in germ cells ^28^. Germline excision of the floxed STOP sequence resulted in permanent activation of the reporter allele, which was transmitted to the progeny independently of Cre expression. Subsequent breeding with wild-type C57BL/6 mice eliminated the Cre transgene, yielding the STAT6-*luc2* line (Figure 1C). Consistent with the physiological activity of the transcription factor ^13,29^, STAT6-*luc2* mice displayed robust and widespread bioluminescent signals throughout the body (Figure 1C and S1A), confirming that germline removal of the STOP cassette effectively restored reporter expression across all tissues.

**Figure 1.**
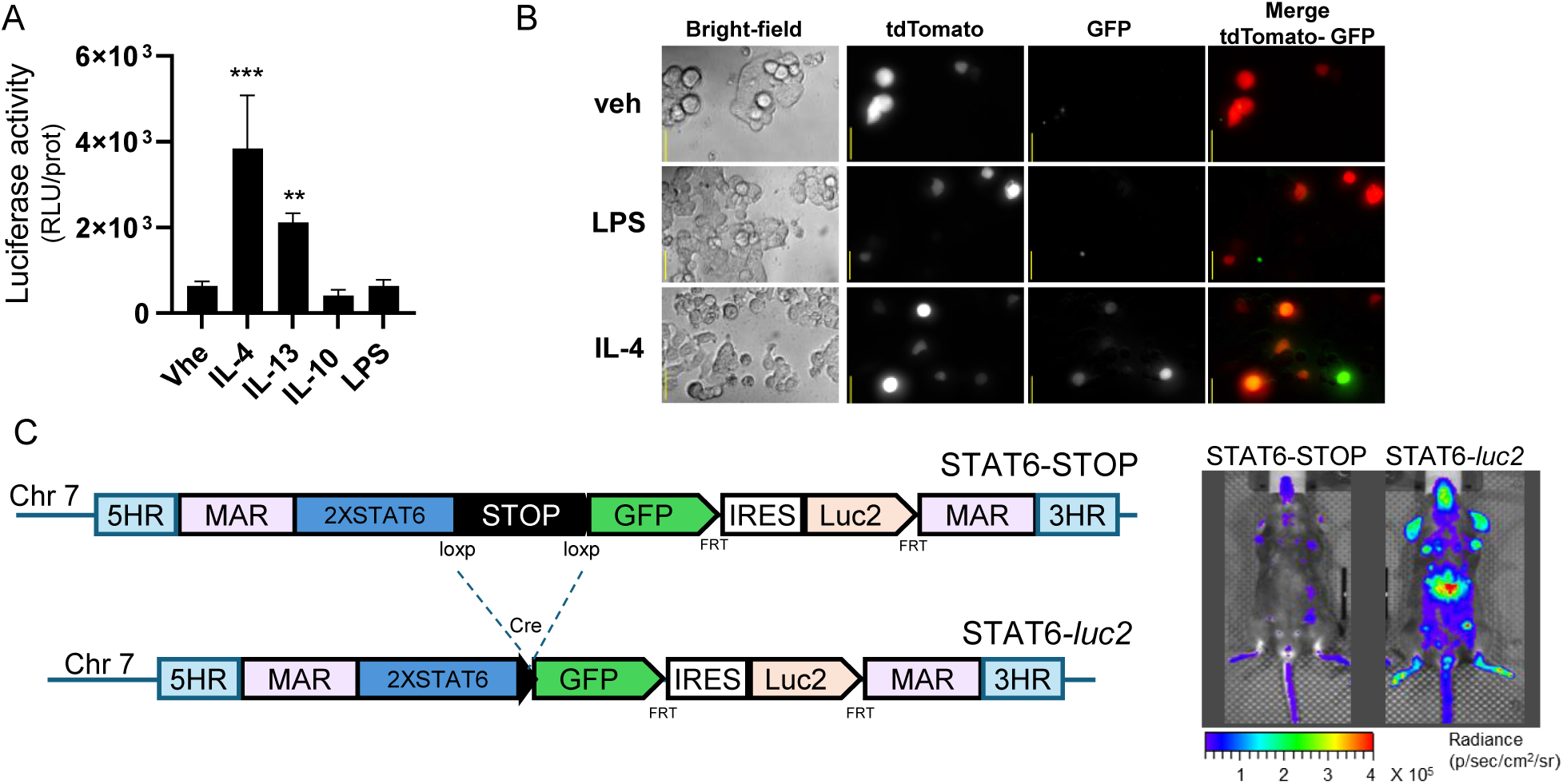
Design and functional analysis of STAT6 reporter systems. **(A)** Luciferase activity was measured in RAW 264.7 cells transiently co-transfected with the *p2xSTAT6*-GFP-*luc2* construct and treated with 20 ng/ml cytokines for 24 h, 2 µg/ml LPS for 6 h, or vehicle (water). Data are mean ± SD (n = 4) of luciferase activity (RLU) normalized to total protein content. ***p < 0.001, **p < 0.01 versus vehicle (one-way ANOVA, Dunnett’s test). **(B)** Representative fluorescence microscopy images showing GFP expression in HT-29 cells transfected with the *p2xSTAT6-GFP-luc2* construct and a constitutively expressing tdTomato plasmid and treated with 20 ng/ml IL-4, 100 ng/ml LPS, or vehicle (water) for 24 h. Scale bar: 50 µm. **(C)** Schematic representation of the STAT6-STOP and STAT6-*luc2* reporter mouse models. The transgene, shown before and after excision of the STOP sequence, was inserted into chromosome 7 (Chr7) of reporter mice by homologous recombination, The mouse line carrying the STOP cassette is referred to as STAT6-STOP. Breeding with *B6.C-Tg(CMV-cre)1Cgn/J mice* (Cre) leads to Cre-mediated excision of the STOP sequence, generating the STAT6-*luc2* line. Abbreviations: 5HR, 3HR:homologous regions for chromosomal integration; MAR: Matrix Attachment Regions; 2XSTAT6: STAT6 promoter; STOP: POLR2 (RNA polymerase II) termination signal; loxP: locus of X-over P1; GFP: green fluorescent protein; *luc2*: optimized firefly luciferase 2; IRES: internal ribosome entry site; Frt: Flp recombination target sites. On the right, representative pseudocolor images of ventral luciferase emission from STAT6-STOP and STAT6-luc2 female mice are shown according to the indicated scale bar.

### Validation of the STAT6-*luc2* reporter mouse

To validate the functionality of the *STAT6-luc2* reporter line, we assessed whether luciferase expression could be induced following activation of STAT6-dependent anti-inflammatory signalling. Mice were intraperitoneally injected with either IL-4 (5 or 50 µg/kg), LPS (50 µg/kg or 1 mg/kg), or vehicle (PBS). Whole-body bioluminescence imaging was performed at 0 h, 5 h and 24 h post-injection to monitor the kinetics of pathway activation (Figure 2A). Bioluminescent imaging revealed a clear, dose-dependent increase in photon emission in the abdominal and pelvic regions 5 h after IL-4 administration, with a moderate signal also detected in the thoracic area (Figure 2B). In contrast, LPS induced only a weak and non–dose-dependent increase in luminescence, confined mainly to the abdominal region and not reaching statistical significance. At 24 h post-administration, luciferase activity returned to baseline levels across all treatment groups, consistent with the transient kinetics typically associated with cytokine-driven STAT6 activation ^13,30^. Comparable responses were observed upon intraperitoneal administration of IL-13, another canonical activator of STAT6 (Figure S1). Notably, under identical experimental conditions, LPS administration (1 mg/kg) elicited a robust inflammatory response in NFκB-*luc2* reporter mice, a reporter mouse model previously generated and validated by our lab ^21^, whereas STAT6-*luc2* mice displayed markedly lower signal intensity (Figure 2A and S1). These findings confirm the pathway specificity of the STAT6 reporter and its selective responsiveness to anti-inflammatory–mediated signalling.

**Figure 2.**
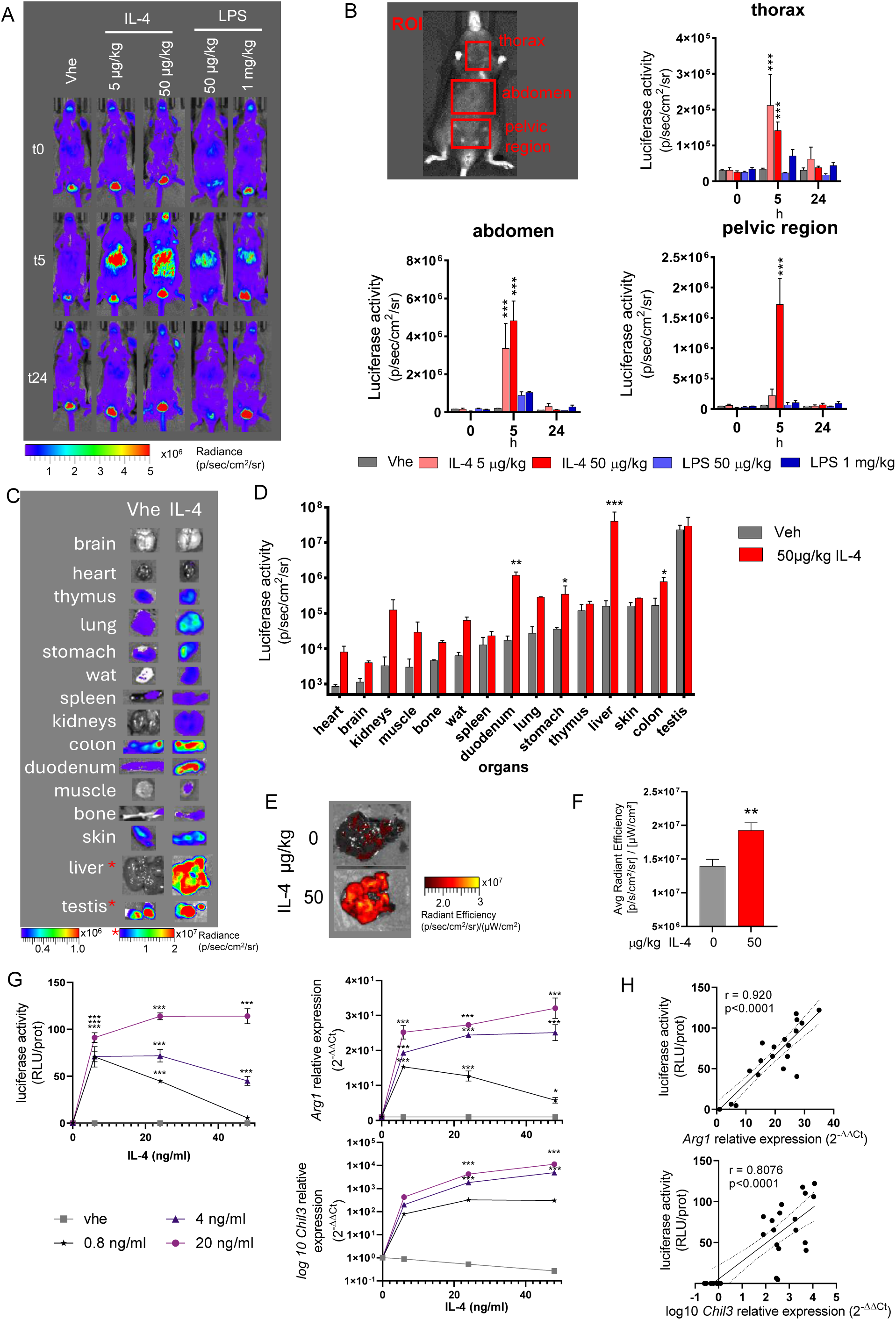
Validation of STAT6-*luc2* reporter mouse. **(A)** Representative *in vivo* bioluminescence imaging of STAT6-*luc2* reporter mice following intraperitoneal injection of IL-4 (5 or 50 µg/kg), LPS (0.05 or 1 mg/kg), or vehicle (PBS). Images were acquired at 0, 5, and 24 h post-treatment. Pseudocolor images represent bioluminescence intensity according to the indicated scale bar. **(B)** Quantification of *in vivo* bioluminescent signal in selected regions of interest (ROIs, red squares). Bars represent mean photon flux (photons/s/cm²/sr) ± SD (n = 3). ***p < 0.001 versus vehicle, (Two-way ANOVA, Dunnett’s test). **(C)** Representative pseudocolor images of bioluminescence emitted from dissected organs collected 5 h after IL-4 (50 µg/kg) or vehicle (PBS) administration and **(D)** quantification of *ex vivo* organ bioluminescence. Bars represent mean photon emission (photons/s/cm²/sr) ± SD (n = 2). *p < 0.05, **p < 0.01, ***p < 0.001 versus vehicle (two-way ANOVA, Sidak’s test). **(E)** Representative pseudocolor images of fluorescence emitted from the liver 5 h after IL-4 (50 µg/kg) or vehicle treatment, and **(F)** quantification of liver fluorescence intensity. Bars represent average radiant efficiency [(p/s/cm²/sr)/(µW/cm²)] ± SD (n = 3). **p < 0.01 versus vehicle, determined by unpaired *t*-test. **(G)** Peritoneal macrophages derived from STAT6-*luc2* mice were treated with IL-4 at different doses (0, 0.8, 4, 20 ng/ml) for 0, 6, 24, or 48 h. At each time point, luciferase expression in protein extracts was quantified and expressed as relative light unit (RLU) per μg of protein ± SEM. At the same time points, relative quantification of target gene transcripts (*Arg1* and *Chil3*) was performed using the 2^−ΔΔCt^ method versus vehicle (*Arg1* above, *Chil3* log10 expression below) ± SEM. ***p < 0.001 versus vehicle (two-way ANOVA, Dunett’s test). **(H)** Correlation between luciferase activity and gene relative expression reported in (G) is shown; Pearson r and p-values for each interpolated curve are reported.

To further characterize tissue-specific reporter activation, *ex vivo* bioluminescence imaging was performed 5 h after IL-4 injection (50 µg/kg). A widespread increase in bioluminescence was observed across multiple organs, with the most pronounced activation detected in the stomach, liver, duodenum, and colon (Figure 2C, 2D) when compared to vehicle-treated mice. Consistent with these results, fluorescent optical imaging of liver tissue revealed detectable GFP signal in IL-4–treated animals, confirming both reporter genes expression (Figure 2E, 2F).

Finally, we sought to determine whether STAT6-*luc2* mice could also serve as a source of primary cells capable of reporting STAT6 activation: peritoneal macrophages were isolated from STAT6-*luc2* reporter mice and stimulated *in vitro* with increasing concentrations of IL-4 (0, 0.8, 4, and 20 ng/ml). Luciferase activity was measured at 0, 6, 24, and 48 h post-stimulation and compared with the expression levels of canonical STAT6 target genes *Arg1* and *Chil3* (also known as *Ym1*)^31^. Luciferase activity closely paralleled *Arg1* expression, displaying a strong positive correlation between reporter induction and *Arg1* mRNA levels (Figure 2G, 2H). A similar trend was observed for *Chil3*, although the correlation followed a logarithmic rather than linear relationship. Comparable results were obtained using bone marrow–derived macrophages (Figure S2).

Collectively, these data demonstrate that the STAT6-*luc2* reporter mouse is a reliable reporter of anti-inflammatory responses and can be used to monitor bioluminescence dynamics *in vivo*, at the tissue level *ex vivo*, and in primary cell cultures.

### Reporter mouse models reveal distinct spatiotemporal modulation of pro- and anti-inflammatory pathways following mRNA vaccination

To elucidate how mRNA-based vaccination modulates pro- and anti-inflammatory responses over time and across distinct anatomical sites, we used two mRNA vaccine formulations encoding the same non-viral antigen derived from an orthologous gene. One vaccine contained an unmodified mRNA sequence (unmod), while the other incorporated N¹-methyl-pseudouridine (mod-m1Ψ) as a chemical modification aimed at reducing innate immune recognition. Both formulations were encapsulated in lipid nanoparticles (LNPs) with identical lipid composition. To assess their immunostimulatory potential, both mRNA formulations were tested *in vitro* using bone marrow–derived dendritic cells (BM-DCs) from wild-type C57BL/6 mice. Cells were treated with 2.5 µg of mod-m1Ψ or unmodified mRNA, their respective LNP controls, or vehicle (water), for 24 and 48 hours. Dendritic cell maturation was assessed by flow cytometric analysis of CD80, CD86, CD40, CCR7, and MHC-II expression, and IL-12 secretion was quantified by ELISA (Figure 3). After 24 hours, only the unmodified mRNA molecule induced a pronounced upregulation of all superficial maturation markers. At 48 hours, both mRNA formulations promoted the upregulation of CD80, CD86, CD40, and CCR7; however, MHC-II expression and IL-12 secretion remained significantly higher in DCs treated with the unmodified vaccine. Neither the LNP nor the vehicle controls induced detectable activation. These data indicate that both mRNA vaccines can promote DC maturation; however, the unmodified sequence elicits a more rapid response and exhibits greater immunostimulatory activity than its modified counterpart.

**Figure 3.**
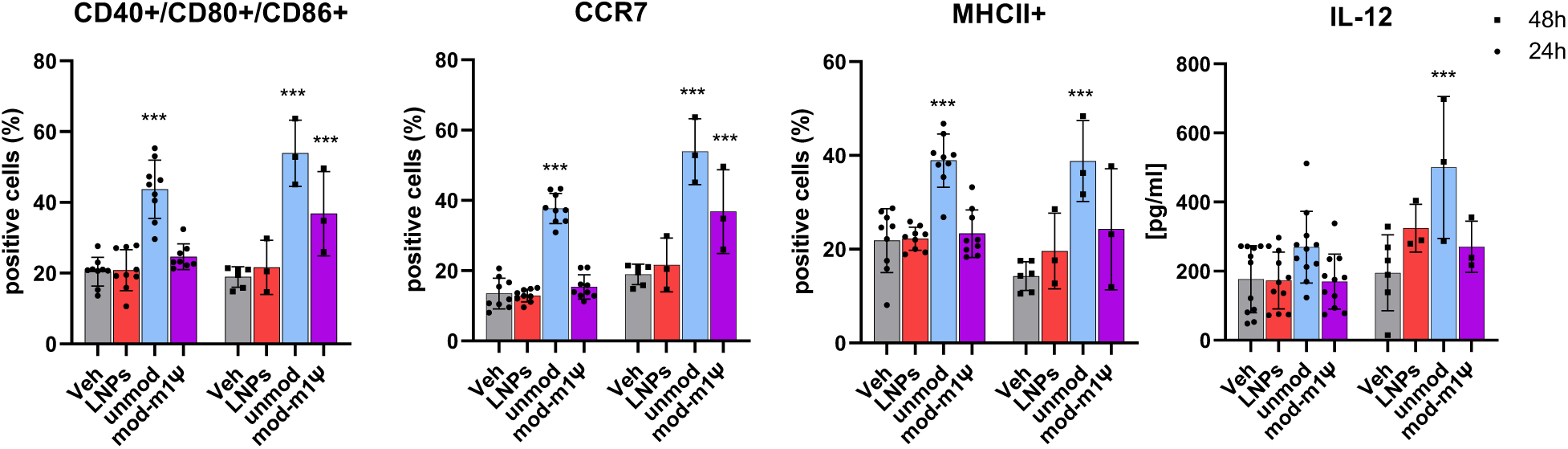
maturation of BM–derived dendritic cells in response to mRNA–LNP treatment. Bone marrow–derived dendritic cells were treated with 2.5 µg of modified or unmodified mRNA encapsulated in LNPs, with an equivalent amount of empty LNPs, or with vehicle (water). Representative histograms show the percentage of CD40-CD80-CD86-CCR7, and MHCII positive cells assessed by flow cytometry, and IL-12 concentrations measured by ELISA ± SEM at 24 or 48 h post treatment (n = 9 at 24 h; n = 3 at 48 h). ***p < 0.001 versus vehicle (two-way ANOVA, Dunnett’s test).

We next moved to *in vivo* study, first assessing the kinetic profile of labelled mRNA–LNPs following tail intravenous administration (i.v). C57BL/6 mice received 10 µg of Cy5.5-labeled, m¹Ψ-modified mRNA, based on the assumption that biodistribution is primarily dictated by the LNP carrier rather than RNA chemistry ^32^. The formulation exhibited rapid systemic distribution, with marked accumulation in the abdominal/pelvic region within 30 minutes, followed by fast clearance (Figure S3), consistent with the expected biodistribution of intravenously administered LNP-based formulations ^33^.

Subsequent, to investigate the spatial and temporal modulation of inflammatory pathways, NFκB-*luc2* and STAT6-*luc2* reporter mice were injected intravenously (10 µg, tail vein) on day 0 with either mod-m1Ψ or unmod mRNA vaccines, or with the corresponding LNP controls, followed by a booster dose on day 14. Bioluminescence imaging of reporter mouse was performed at 0, 5 hours, 1 day, and 6 days after injections, and again on day 29 (14 days after the booster) to monitor reporter activation across different anatomical regions (Figure 4).

**Figure 4.**
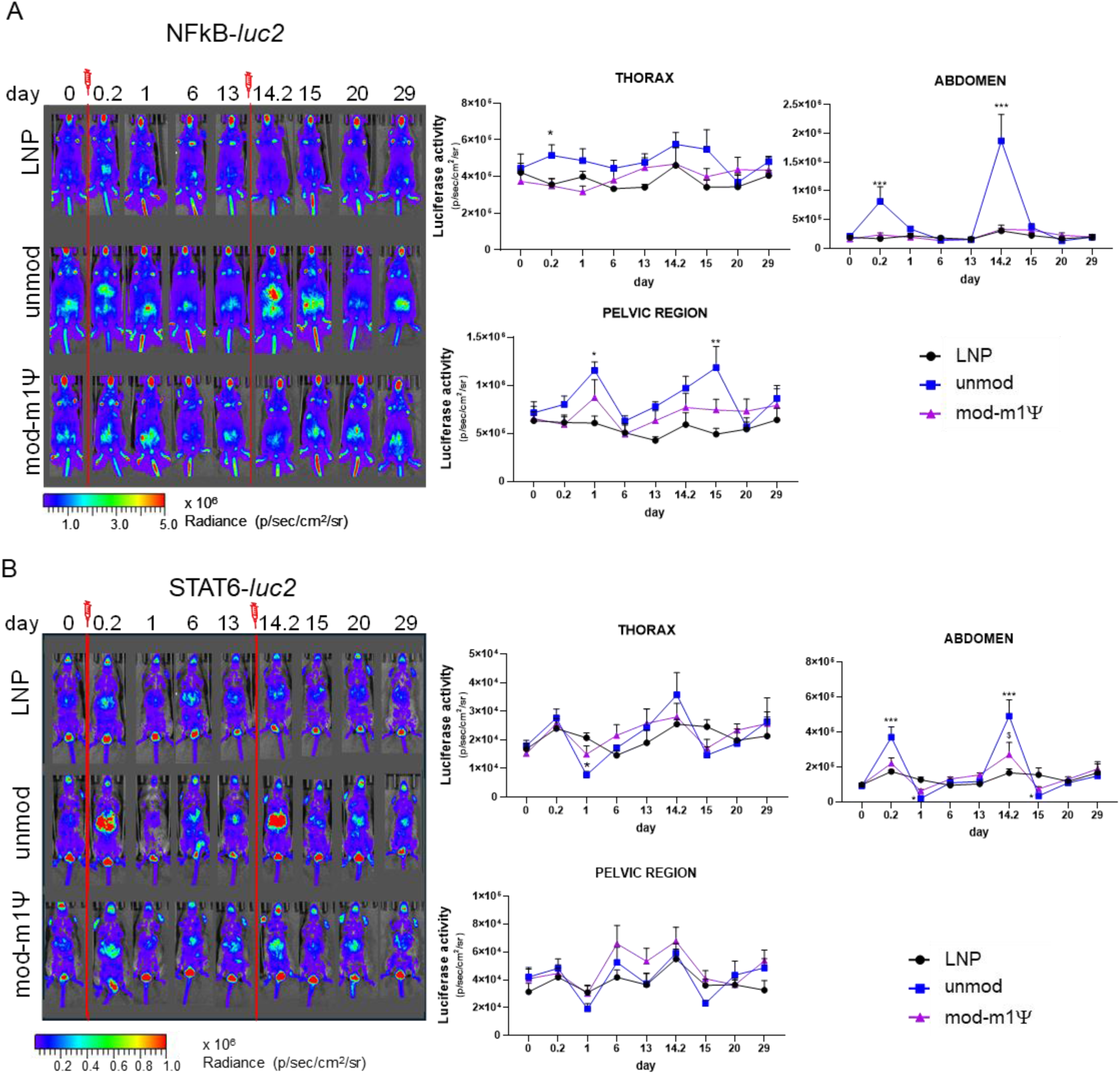
*In vivo* proinflammatory and anti-inflammatory mRNA vaccine responses in reporter mice. **(A)** NFκB-*luc2* mice or **(B)** STAT6-*luc2* were injected intravenously (10 µg/kg) with unmodified mRNA vaccine (unmod), modified mRNA vaccine (mod-m1Ψ), or the corresponding amount of LNPs on days 0 and 14. Representative longitudinal *in vivo* bioluminescence images (0, 0.2, 1, 6, 13, 14.2, 15, 20 and 29 day) are shown on the left, displayed using pseudocolors according to the scale bar. Bioluminescence was quantified in selected regions of interest as defined in Figure 2. Bar graphs represent mean photon emission (photons/s/cm²/sr) ± SEM (n = 6). *p < 0.05, **p < 0.01, ***p < 0.001 versus LNP (two-way ANOVA, Dunnett’s test).

In NFκB-*luc2* mice (Figure 4A), the unmod vaccine induced a robust and rapid bioluminescent signal in the abdominal region as early as 5 hours after the first and booster administrations, followed by prominent activation in the pelvic region at 1 day after both administrations. A mild but sustained increase in signal in the thoracic region was observed in mice receiving the unmodified vaccine from 5 hours (day 0.2). In contrast, the mod-m1Ψ vaccine elicited a modest increase in the pelvic signal on day 1 and again from days 13 to 29, although these changes did not reach statistical significance. In STAT6-*luc2* mice (Figure 4B), both mRNA vaccines induced in the abdominal region a detectable activation of the anti-inflammatory pathway 5 hours after the booster injection, and the unmod mRNA also elicited activation 5 hours after the primary dose; the signal was consistently stronger in mice receiving the unmod vaccine. However, a pronounced reduction in signal intensity was observed 24 hours post-administration across all regions, with the most significant decreases in the thoracic and abdominal areas for the unmod sequence, indicating a transient activation followed by downregulation of this pathway.

Taken together, these results provide the first evidence of spatiotemporal activation of pro- and anti-inflammatory signalling pathways following systemic mRNA vaccination. Shortly after administration, the earliest and most prominent responses were observed in abdominal area (predominantly containing the liver), for both NF-κB and STAT6 signalling. In this region, NF-κB activation was stronger and more sustained in response to unmodified mRNA, returning to baseline levels approximately 24 h after administration. In contrast, STAT6 activation was detected following both vaccines. Notably, at 24 h post-administration, STAT6 activity decreased below baseline levels, supporting the hypothesis of an active shutdown of STAT6-dependent transcription following an initial activation phase. Given that this reduction was observed across nearly all anatomical regions, it suggests a generalized systemic response. Overall, these data indicate that abdominal organs exhibit pronounced STAT6 and NF-κB modulation during the early phase of systemic mRNA vaccine administration.

### *Ex vivo* assay to study organ-specific activation

To obtain a comprehensive view of the organ-specific activation profile, bioluminescence imaging was performed on dissected organs both 5 hours after i.v. vaccine administration and at the end of the experiment (day 29) in reporter mice, following the experimental setup previously described. This analysis enabled the identification of organs primarily involved in STAT6 and NF-kB modulation. At the same time-point, cytokine production profiles were assessed in plasma, to further characterize the immune responses elicited by the two vaccine formulations, while antigen-specific antibody titres were quantified on day 29. A cohort of vehicle-treated animals (glucose solution) was included at the 5-hour time point to better delineate the contribution of the LNP component.

At 5 hours post-administration (Figure 5A), NF-κB activation was observed in the thymus and liver of mice receiving the unmodified sequence, with a general upward trend across multiple organs. In contrast, mice treated with the modified sequence or with LNPs alone exhibited activation levels comparable to vehicle-treated controls, except for a mild, non–statistically significant activation detected in the liver of the mod-m1Ψ group. Regarding STAT6 activation, the unmodified sequence induced a high level of activation in the liver, whereas LNP and mod-m1Ψ elicited comparable bioluminescence signals, both higher than those observed in vehicle-treated mice. In contrast, the mod-m1Ψ mRNA predominantly triggered activation in the thymus.

**Figure 5.**
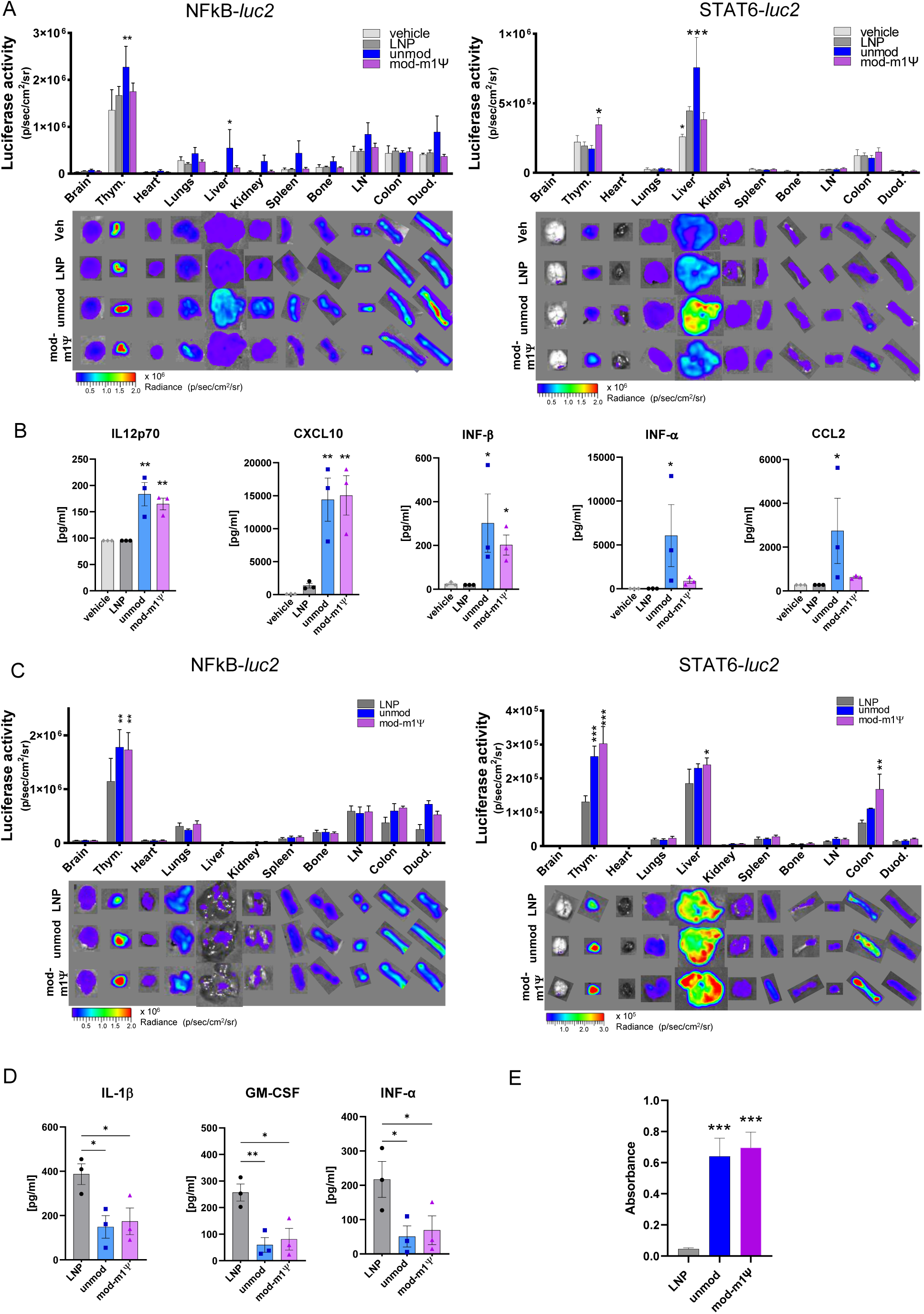
*Ex vivo* analysis of pro- and anti-inflammatory responses at 5 hours and 29 days, plasma cytokine levels, and antigen-specific antibodies. **(A, C)** Representative *ex vivo* bioluminescence images of organs dissected from NFκB*-luc2* and STAT6*-luc2* reporter mice at (A) 5 hours or (C) 29 days after treatment. Signals are shown using pseudocolors according to the scale bar. Quantification of organ bioluminescence is shown in the bar graphs, representing mean photon emission (photons/s/cm²/sr) ± SEM (n = 6). *p < 0.05, **p < 0.01, \*\**p < 0.001 versus* LNP (two-way ANOVA, Dunnett’s test).Thym: thymus, LN: inguinal lymph nodes, Dued.: duodenum. **(B, D)** Cytokine levels in plasma were measured (B) 5 hours after administration or (D) at day 29. Cytokine concentrations are presented as bar graphs ± SEM (n = 3). *p < 0.05, \**p < 0.01 versus* LNP (one-way ANOVA, Dunnett’s test). **(E)** Antigen-specific antibodies were measured in plasma at day 29 by ELISA. Bar graphs represent mean absorbance ± SEM (n = 12). \*\**p < 0.001* versus LNP (two-way ANOVA, Dunnett’s test).

Cytokine profiling in the plasma at 5 hours after injection revealed upregulation of IL-12p70, CXCL10, and IFN-β in both vaccine groups relative to control and LNP-treated mice, whereas CCL2 and IFN-α were selectively increased in animals administered the unmodified mRNA, while no statistical significant modulation was observed for IL-1β, IL-10, INF-γ, CXCL1 (Figure 5B and S4). These data indicate that both vaccines induced distinct immunostimulatory signatures primarily attributable to the mRNA chemistry rather than to the LNP formulation.

At day 29 (Figure 5C), bioluminescence imaging demonstrated sustained activation of NF-κB and STAT6 in the thymus of both vaccine groups. Moreover, STAT6 activation persisted in the liver and colon of mice treated with the modified sequence. The absence of unspecific upregulation of either NF-κB or STAT6 pathways in other organs suggests that the inflammatory response remained moderate and spatially confined at this time point. Notably, cytokine analysis revealed downregulation of IL-1β, GM-CSF, and IFN-α in both vaccine groups consistent with resolution of the early proinflammatory response, while no statistical differences were detected for INF-γ, CXCL1, CCL2, IL12P70, IL-10, IL-6 (Figure 5D and S4).

Finally, analysis of anti-antigen antibodies in plasma at day 29 showed that both vaccine formulations elicited detectable antigen-specific responses, with no significant differences in antibody titres between the two groups (Figure 5E).

### Identification of important patter of immunogenic potential

Given that the two vaccine formulations exhibited distinct NF-κB and STAT6 activation profiles while ultimately inducing comparable antibody responses, we hypothesized that correlating bioluminescence data with antibody production could help identify key time points, organs, and anatomical regions relevant for effective immunization. This approach aimed to define which organ-specific and temporal activation profiles of STAT6 and NF-κB are most conducive to robust antibody production.

Initially, we correlated bioluminescence intensity measured at day 29 in dissected organs (Figure 5C) from LNP-, modified-, and unmodified-treated mice with antigen-specific antibody levels (Figure 5E). As shown in Figure 6A, STAT6 activation exhibited a strong positive correlation with antibody titres in both the thymus, colon and duodenum whereas NF-κB activity did not display any consistent association with antibody production at any time point.

**Figure 6.**
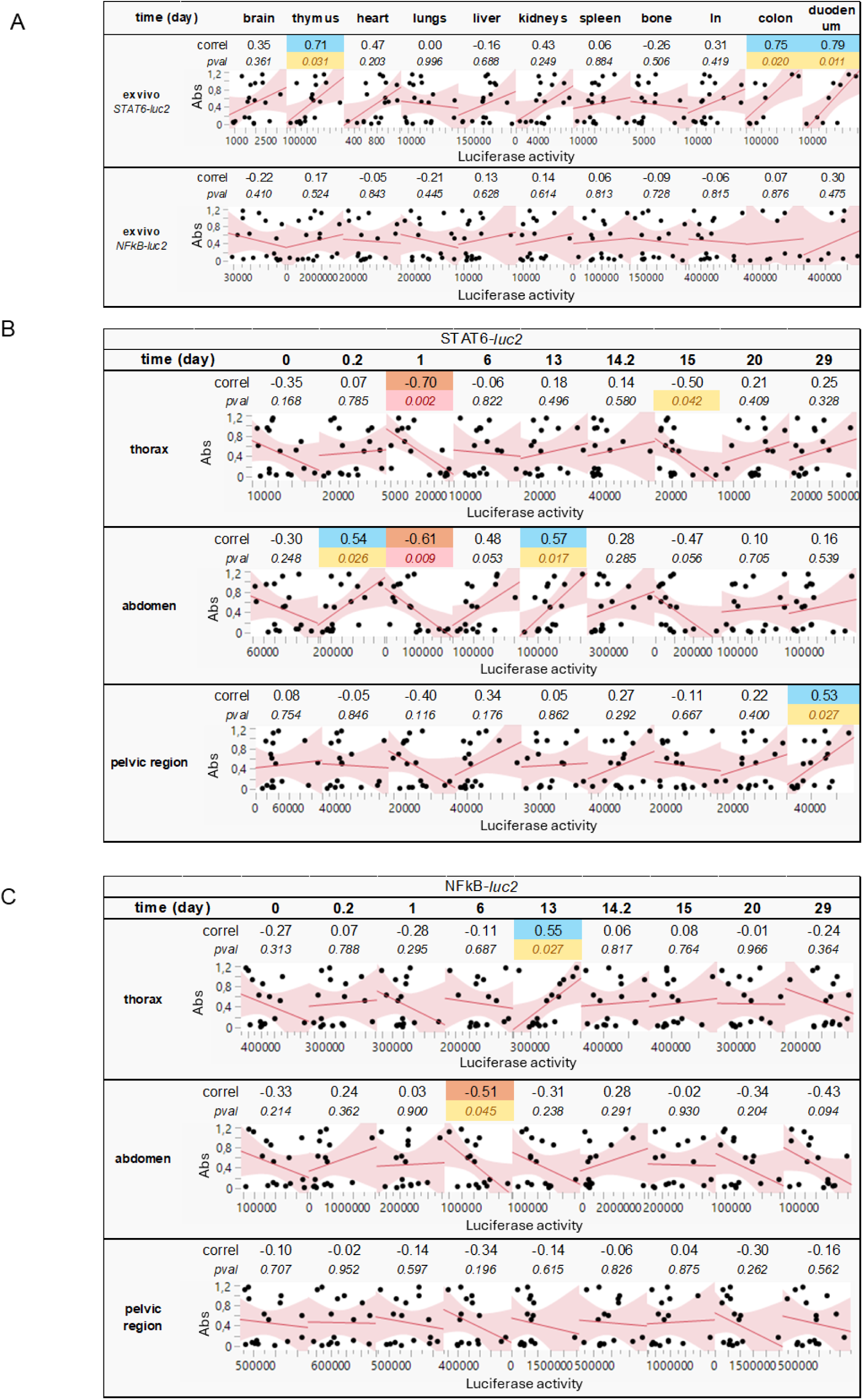
Correlations between antigen-specific antibodies and bioluminescence signals. **(A)** Correlations between *ex vivo* bioluminescence of dissected organs at day 29 (abscissa, luciferase activity (photons/s/cm²/sr)) and antigen-specific antibody absorbance (ordinate, Abs) values are shown, with Pearson’s correlation coefficients (r; correl.) and p-values (pval) for each interpolated curve. ln: lymph nodes. **(B, C)** Correlation analyses between *in vivo* bioluminescence signals in selected regions at defined time points (abscissa, luciferase activity (photons/s/cm²/sr)) and antigen-specific antibody absorbance in plasma at day 29 (ordinate, Abs) are shown for (B) *STAT6-luc2* mice and (C) *NFκB-luc2* mice. Pearson’s r (correl) and corresponding p-values (pval) are reported for each interpolated curve.

We next investigated whether the *in vivo* bioluminescence dynamics were associated with the vaccine’s capacity to induce an effective humoral response. To this end, bioluminescent signals detected in specific anatomical regions at different time points were correlated with plasma antibody levels measured at day 29. As shown in Figure 6B and 6C, several time points and anatomical regions displayed statistically significant correlations between bioluminescence intensity and antibody titres. Notably, the early STAT6-mediated response appeared to be the most predictive of immunogenic potential (Figure 6B). Specifically, abdominal bioluminescence at 5 hours, day 13, and in the pelvic region at day 29 post-immunization exhibited strong positive correlations with antibody levels at day 29. Conversely, inverse correlations were observed for STAT6 signals detected at 1-day post-immunization in the abdominal and thoracic regions, as well as at day 15 in the thoracic area.

Regarding NF-κB activity, a positive correlation was observed in the thoracic region at day 13, whereas an inverse correlation was found in the abdominal region at day 6 (Figure 6C).

Together, these findings suggest that early STAT6 activation is more predictive of a successful immune response than NF-κB activation. In contrast, the subsequent attenuation of STAT6 activity in abdominal and thoracic organs appears to be required for optimal immunization. Moreover, the presence of distinct “waves” of STAT6- and NF-κB–mediated activation across different anatomical regions supports the notion of a temporally and spatially coordinated immune regulation, underscoring the complex interplay of these signalling pathways during vaccine-induced immunity.

## Discussion

In this study, we report the first validation of a STAT6 reporter mouse capable of real-time, whole-body monitoring of STAT6-dependent immunomodulatory activity. This model enables both dynamic *in vivo* imaging and organ-specific *ex vivo* analyses. Efficient excision of the STOP cassette confirms the functionality of the Flox system, indicating that crossing STAT6-STOP reporter mice with tissue-specific Cre drivers could facilitate compartment-restricted investigation of STAT6 biology. STAT6-*luc2* reporter mice faithfully detect the expected anti-inflammatory responses following administration of cytokines such as IL-4 and IL-13. Notably, the responses exhibit spatial and dose-dependent differences, with a prominent abdominal bioluminescence signal primarily originating from the liver—an organ characterized by high *Arg1* expression ^29,34^ and well-established sensitivity to STAT6-mediated transcription ^35^. These features establish the model as a powerful platform for *in vivo* investigation of STAT6 signalling. Noteworthy, this system provides complementary insights to the pro-inflammatory responses captured by NFκB-*luc2* reporter mice ^21^ When used in combination, these dual-reporter models offer a comprehensive and high-resolution physiological view of immune activation, representing a valuable toolkit for studying the spatial and temporal regulation of immunity *in vivo*.

Applying these reporter mice to LNP-encapsulated mRNA vaccination enabled a spatially and temporally resolved comparison of immune modulation by two vaccine formulations encoding the same antigen but differing in RNA chemistry. As expected, unmodified mRNA elicited stronger early innate activation than its N¹-methyl-pseudouridine (m¹Ψ)-modified counterpart, reflected in heightened NF-κB signalling. This is consistent with the reduced TLR7/8 and RIG-I stimulation by m¹Ψ-containing transcripts ^22^ and corroborated by plasmatic cytokine profiling at 5 hours post-administration, which revealed induction of IFN-α and CCL2 specifically by unmodified mRNA. In fact, within hours of intravenous administration, NF-κB activation was pronounced in the liver and thymus for the unmod vaccine, concurrently, STAT6 activation was observed in the abdomen and liver, whereas the modified formulation induced STAT6 signalling primarily in the thymus. Interestingly, LNPs alone were sufficient to trigger STAT6 responses at 5 hours post administration. The hepatic signal is consistent with the LNP pharmacokinetics and preferential hepatic accumulation ^36^.

Strategically located at the confluence of arterial and venous blood, the liver continuously monitors both systemic and gut-derived molecules. It contains a dense network of immune cells—including Kupffer cells (80–90% of all tissue macrophages), DCs, and natural killer cells—expressing diverse immune receptors ^37^. Despite this, the liver favours immunological tolerance, preventing unnecessary responses to harmless stimuli. Under non-threatening conditions, such as gut-derived LPS without inflammation, Kupffer cells remain tolerogenic and limit T cell activation; however, additional pathogen signals or inflammatory cytokines can reprogram them into potent antigen-presenting cells^37^. In our system, the liver exhibits a strong and early response, emphasizing the need for detailed hepatic analysis during immunization, as it may play a central role in determining whether an immune response is initiated.

By 24 hours, a new wave of NF-κB activation emerged in the pelvic region for both vaccines, associated with a systemic reduction in STAT6 signalling. The timing is consistent with the 6–24-hour window required for DCs migration and initiation of antigen presentation ^38^ and this pattern likely reflects activation of lymph nodes during the onset of T-cell priming ^11,12^. The reduction of STAT6 signalling by 24 hours is consistent with its well-described transient kinetics. Early responses to vaccination are dominated by Th1-polarizing cytokines and type I interferons, which inhibit STAT6 phosphorylation, suppress IL-4 production, and limit Th2 differentiation ^2–4^. Prolonged STAT6 activity would risk undesirable Th2 polarization, IgE class switching, and pro-fibrotic gene expression; therefore, the reporter probably highlighting the feedback mechanisms actively terminate STAT6 signalling^39,40^. The absence of sustained STAT6 activity is therefore mechanistically consistent with effective immunization and likely supports immunological protection.

Notably, at 29 days, NF-κB and STAT6 activity persisted in the thymus for both vaccines, consistent with ongoing processes such as thymocyte proliferation and the balance between maturation and apoptosis ^2–4^ . The failure to detect significant changes in luciferase expression in lymph nodes and spleen, where mechanistic effects would be anticipated, may reflect kinetic differences in pathway activation or limited statistical power due to low signal intensity, necessitating increased sample sizes. Interestingly, for the modified vaccine, STAT6 activation was also observed at 29 days in the liver and colon, suggesting organ-specific long-term immune modulation beyond the thymus.

Only few studies have examined the immunological impact of LNP-based vaccines on the liver, our observations align with Pongma et al. ^41^, who demonstrated that 7 day after mRNA/LNPs vaccination, liver-associated macrophages undergo alteration in gene expression and epigenetic profiles, highlighting the liver’s active participation in immune responses rather than acting as a passive nanoparticle depot. The lack of systemic NF-κB activation aligns with the observed reduction in pro-inflammatory plasmatic cytokines—including IL-1β, IFN-α, and GM-CSF—at 29 days, suggesting that these vaccines do not elicit prolonged inflammatory responses that could be potentially detrimental.

Integration of imaging bioluminescent data with antibody measurements revealed that early pathway dynamics were predictive of immunogenicity. Among all time points, the most informative and predictive were the early (5 h) STAT6 activation in the abdomen, followed by rapid deregulation in the abdomen and thorax by 24 hours, signals that correlate with robust antibody responses. This identifies STAT6 as a useful early biomarker of effective vaccine-induced immunity. In contrast, strong early NF-κB activation did not correlate with immunogenic antibody production, consistent with previous reports suggesting that excessive inflammation may compromise optimal adaptive responses ^42^. Together, these findings demonstrate that NF-κB and STAT6 reporter mice provide a high-resolution platform for assessing vaccine performance and dissecting early immunoregulatory pathways. Because predictive signatures emerge within the first 24 hours, this strategy system could be further developed to enable rapid *in viv*o screening of candidate vaccines, allowing prioritization of leads that recapitulate the transcription factor dynamics of highly immunogenic vaccines—without waiting for antibody readouts—and thereby accelerating vaccine development.

## Conclusions

In summary, we present and validate complementary STAT6 and NF-κB reporter mice that enable real-time, organ-resolved monitoring of key regulatory pathways shaping early immune responses. Using these tools, we demonstrates that LNP-based mRNA vaccination elicits highly coordinated, yet chemistry-dependent, immune dynamics. This enables, the first time, detailed analysis of the spatiotemporal patterns of STAT6 and NF-κB signalling, with the liver emerging as a dominant early hub for both pathways. These findings suggest that hepatic immune cross-talk should be studied in detail, as it may represent a critical determinant of downstream adaptive immunity during LNP-mediated vaccination. In addition, our data identify transient early STAT6 activation—rather than strong NF-κB–driven inflammation—as a positive predictor of antibody responses, highlighting the importance of balanced innate signalling in effective immunization. These insights support the use of STAT6 and NF-κB reporter mice as rapid *in vivo* screening platforms to identify promising vaccine formulations within the first hours, substantially accelerating preclinical evaluation.

## Materials and Methods

### Plasmids construction

The plasmid *p2xSTAT6-GFP-luc2* was generated by inserting the promoter region from *p2xSTAT6-Luc2P* (a gift from Axel Nohturfft; Addgene plasmid #35486, RRID:Addgene 35486) upstream of the *loxP-STOP-loxP-GFP-FRT-IRES-luc2-FRT-polyA* cassette using the *XhoI* and *MluI* restriction sites. The floxed STOP cassette was then excised using Cre recombinase (New England Biolabs, M0298S) according to the manufacturer’s protocol prior to performing cellular transfection assays. The FRT sites could be used to excise *luc2* through FLP recombinase. To generate the targeting vector, the resulting *2xSTAT6-loxP-STOP-loxP-GFP-flox-IRES-luc2-flox* cassette was cloned into the targeting backbone via site-directed recombination (VectorBuilder). In the final construct, the transgene was flanked by matrix attachment regions (MARs) and by homologous arms corresponding to chromosome 7, locus 21 ^21^.

### Generation of STAT6-STOP and STAT6-luc2 reporter mouse

The targeting vector was linearized with *NotI* and transferred into 129/Sv embryonic stem cells by electroporation, using 35 μg of DNA for each 15 million cells (Core Facility for Conditional Mutagenesis, DIBIT San Raffaele). Positive clones were selected with puromycin (1 μg/ml), and one hundred resistant clones were screened for homologous recombination by PCR. A positive clone was injected into C57BL/6NCrL blastocysts and transferred to pseudo-pregnant CD1 females. The resulting chimeric male mice, with approximately 80-90% chimerism, were bred with wild-type C57BL/6J female mice to produce F1 transgenic mice named STAT6-STOP. This line was then crossed with B6.C-Tg(CMV-cre)1Cgn/J mice to obtain STAT6-*luc2* reporter mice.

#### Immortalized cell cultures and transient transfections

RAW 264.7 and HT-29 cell lines were obtained from the American Type Culture Collection (ATCC) and cultured in cultured in Dulbecco’s Modified Eagle Medium (DMEM; Gibco, Cat.11965-092) supplemented with 10% fetal bovine serum (FBS; Gibco, Cat. A5209502), 1 mM Sodium Pyruvate (Gibco, Cat.11360070) and 1% Antibiotic-Antimycotic (Gibco, Cat. 15240-062), and maintained at 37 °C in a humidified atmosphere containing 5% CO_2_. Both cell lines were transfected with *p2xSTAT6-GFP-luc2* plasmid. For transfection, 50 000 Raw 264.7 cells were seeded in a 24-well plate and 60 000 HT-29 cells were seeded in a 48-well plate, cultured overnight prior to transfection. Transfection of RAW 264.7 cells were performed using FuGENE HD Transfection Reagent (Promega, Cat. 2311) with a DNA:Fugene ratio of 2:6 (µg:µl) for each well, for HT-29 was performed using Lipofectamine LTX Transfection Reagent (Thermo Fisher Scientific, Cat. 15338100) with a DNA:Lipofectamine ratio of 1:4.8 (µg:µl) following the manufacturer’s instructions. HT-29 cells were co-transfected with *the pCDH-EF1-Luc2-P2A-tdTomato* (a gift from Kazuhiro Oka - Addgene plasmid #72486, 10% of total DNA). Transfected cell lines were treated with IL-4, IL-13, IL-10 (PeproTech, Cat. 200-04, 200-13, 200-10, respectively), for 24h or LPS (Merck, Cat. L2630) for 6h or 24h. Fluorescence images of live transiently transfected cells were acquired using an Axiovert 200M microscope with dedicated software (AxioVision Rel 4.9, Zeiss) at a magnification of ×20.

#### Bone-marrow derived and peritoneal macrophages isolation and growth

Mouse peritoneal macrophages were isolated by peritoneal lavage using 5 ml of pre-chilled 0.9% NaCl injected into the peritoneal cavity with a 21G needle. Recovered cells were centrifuged at 1500 rpm for 5 min and subjected to red blood cell lysis with ACK buffer (0.15 M NH_4_Cl, 1 mM KHCO_3_, 0.1 mM EDTA; pH 7.3) for 5 min at 4°C. Cells were then seeded at 1 × 10⁶ cells/well in 24-well plates in RPMI+GlutaMAX (Thermo Fisher Scientific, Cat. 61870036) supplemented with 10% endotoxin-free FBS (Gibco, Cat. A5209502), 1% penicillin/streptomycin (Gibco, Cat. 15240-062), and sodium pyruvate (Gibco, Cat. 11360070). After 45 min, non-adherent cells were removed by extensive PBS washing, and adherent macrophages were maintained in RPMI containing 10% FBS. Treatments were performed the following day.

Bone marrow–derived macrophages (BMDMs) were generated by flushing bone marrow from tibia and femur with DMEM+GlutaMAX (Gibco, Cat. 11965-092) using a 21G needle. Cells were centrifuged at 1250 rpm for 8 min, seeded in T75 flasks in DMEM+GlutaMAX supplemented with 10% endotoxin-free FBS, 1% penicillin/streptomycin, and 1% sodium pyruvate, and incubated overnight. The following day, cells were collected, centrifuged, and reseeded at 3 × 10⁶ cells/dish for differentiation over 6 days in DMEM+GlutaMAX containing 20% endotoxin-free FBS, 30% L929 cell–conditioned medium (ATCC), 1% penicillin/streptomycin, and 1% sodium pyruvate. After differentiation, BMDMs were detached using Accutase (Merck-Millipore, Cat. SCR005) and plated in 24-well plates at 4.5 × 10⁵ cells/well. After 45 min, cells were extensively washed with PBS and maintained in DMEM containing 10% FBS. Treatments were performed the following day.

#### Animal treatments and in vivo and ex vivo imaging

All animal experimentation was carried out in accordance with the Animal Research: Reporting of In Vivo Experiments (ARRIVE) guidelines and the European Guidelines for Animal Care. The animal study protocols were approved by: the Italian Ministry of Research and University and by the Finnish National Animal Experiment Board (Hankelupalautakunta – ELLA) (permission numbers: 417/2018, 416/2022, 904/2023, 105/2024). Reporting of In Vivo Experiments (ARRIVE) guidelines and the European Guidelines for Animal Care. Male mice of 3-6 months were used unless otherwise specified. The animals were provided ad libitum access to food and housed in individually ventilated plastic cages at a temperature range of 22-25°C with a relative humidity of 50% ± 10%. The housing environment followed an automatic cycle of 12 h of light and 12 h of darkness. To minimize any circadian influence, both the treated and control groups were analysed at the same time, and the reference point (t0) for the start of the experiments was set in the morning (between 9:00 and 10:00 a.m). For *in vivo* validation, an intraperitoneal injection of IL-4, IL-13 (PeproTech, Cat. 214-14 and 210-13) or LPS (Merck, Cat. L2630) was performed in a saline-based solution. For vaccine experiment a tail vain injection was performed with 10 µg of mRNA or corresponding amount of empty LNP or vehicle (glucose-based solution). For semi-quantitative bioluminescent photon-emission analysis, we followed established protocols ^43^. Mice received a subcutaneous injection of luciferin (Promega, Cat. P1041; 80 mg/kg) 10 minutes prior to imaging. Animals were anesthetized with isoflurane and maintained under anaesthesia throughout each 2-minute optical imaging session. Bioluminescent imaging was performed using a charge-coupled device (CCD) system (IVIS SpectrumCT, PerkinElmer). Following the final *in vivo* acquisition, mice were euthanized via cervical dislocation. Organs were immediately subjected to a 2-minute *ex vivo* imaging session and then rapidly frozen at –80 °C for subsequent analyses. Photon emission in defined regions was quantified using Living Image Software v4.2 (PerkinElmer).

For biodistribution analyses with the fluorescent vaccine, mice were acquired using IVIS SpectrumCT (PerkinElmer), at different time points: after 2, 10, 30 minutes, and 6, 24, 48h after tail vein injection of the fluorescent vaccine. At the end of the experiments, images were scaled and analysed by applying the same regions of interest (ROI), using the Living Image 4.7.4 Software (Perkin Elmer). Fluorescent signals were expressed as average radiance efficiency ([photons/second/square centimeter/steradian]/[[µW/square centimeter]) which is a calibrated measurement of photon emission.

#### Vaccine production: plasmid DNA design, expansion, and template linearization

Plasmid DNA for in vitro transcription (IVT) was designed to include an optimized T7 promoter compatible with CleanCap AG(3’OMe) co-transcriptional capping, a human HBB-derived 5′ untranslated region (5′UTR), and a coding sequence (CDS) consisting of a mouse Ig κ-chain signal peptide fused in-frame to the antigen of interest. The 3′ region of the construct contained two tandem copies of the human HBB-derived 3′UTR (HBB2x), followed by a 100-nt poly(A) tail positioned immediately upstream of a unique *HindIII* restriction site within the pUC57 backbone to enable template linearization. The plasmid was synthesized by GenScript and sequence-verified. The plasmid was transformed into *Escherichia coli Stbl3* chemically competent cells (Invitrogen, Cat. C737303) by heat-shock transformation. Briefly, plasmid DNA was added to competent cells and incubated on ice for 30 min, followed by heat shock at 42 °C for 45 s and immediate transfer to ice for 5 min. Pre-warmed Luria–Bertani broth was added, and cells were recovered by shaking at 37 °C for 1 h before plating on LB agar plates containing ampicillin (100 µg/ml). Plates were incubated overnight at 37 °C. Single colonies were inoculated into LB broth supplemented with ampicillin (100 µg/ml) and grown with shaking at 37 °C for plasmid expansion. Plasmid DNA was isolated using a Macherey-Nagel NucleoBond Xtra Maxi kit (Cat. 740424.10) according to the manufacturer’s instructions. Purified plasmid DNA was quantified spectrophotometrically using a NanoPhotometer NP80 (Implen).

For IVT template preparation, plasmid DNA was linearized by digestion with *HindIII* (New England Biolabs, Cat. R3104L) at a ratio of 5 U enzyme per µg of DNA. Digestions were performed using plasmid DNA at a concentration of 100 ng/µl and incubated for 1.5 h at 37 °C. Linearized DNA was purified by phenol–chloroform–isoamyl alcohol extraction (saturated with 10 mM Tris-HCl, pH 8.0, and 1 mM EDTA; Merck, Cat. P3803) by adding an equal volume of organic solvent, mixing thoroughly, and centrifuging at 14,000 × g for 10 min to separate phases. The aqueous phase was recovered, and DNA was precipitated by addition of 1/10 volume of 3 M sodium acetate (pH 5.2) and isopropanol, followed by overnight incubation at −20 °C. DNA pellets were washed with 70% ethanol, air-dried, and resuspended in nuclease-free water. DNA concentration was re-quantified, and complete linearization was confirmed by agarose gel electrophoresis on a 1% agarose gel run at 100 V using a 1 kb DNA ladder (New England Biolabs, Cat. N3232S).

#### Vaccine production: in vitro transcription

Unmodified mRNA, N¹-methyl-pseudouridine–modified mRNA (m¹Ψ-UTP), and Cy5.5-labeled m¹Ψ-modified mRNA were synthesized by in vitro transcription (IVT) using the MEGAscript T7 kit (Invitrogen, Cat. AMB13345), following the manufacturer’s instructions. IVT reactions were performed using linearized plasmid DNA templates at a final concentration of 50 ng/µL. For unmodified transcripts, reactions contained standard ribonucleotide triphosphates (7.5 mM each) and CleanCap AG (3′-O-Me) (TriLink BioTechnologies, Cat. N-7413) at 5 mM. For modified mRNA, UTP was replaced with N¹-methyl-pseudouridine triphosphate (m¹Ψ-UTP; TriLink BioTechnologies). Fluorescently labelled mRNA was generated by supplementing the nucleotide mix with Cy5-UTP (MedChemExpress, Cat. HY-150145) at a 1:5 Cy5-UTP:UTP ratio. Add the linearized pDNA template concentration (50ng/µL).

All IVT reactions were incubated at 37 °C for 3 h and subsequently treated with TURBO DNase for 15 min at 37 °C to remove residual plasmid DNA. mRNAs were then purified by LiCl precipitation by adding 7.5 M LiCl to a final concentration of 2.5 M, followed by overnight incubation at −20 °C. RNA pellets were recovered by centrifugation at 14,000 × g for 20 min, washed once with 70% ethanol, briefly air-dried, and resuspended in nuclease-free water. RNA concentration and purity were measured using a NanoPhotometer NP80 (Implen), and transcript integrity was assessed by electrophoresis on a 1% agarose gel in 1× TAE buffer, run at 90 V for 1 h with an RNA ladder (New England Biolabs, Cat. N0362S).

### *Vaccine production: m*icrofluidic mRNA-LNP formulation

mRNA–LNPs were formulated using the Sunshine microfluidic system (Unchained Labs) equipped with a 190XT mixer. Lipids were dissolved in ethanol at molar ratios of ALC-0315 (47.4%), cholesterol (40.9%), DSPC (10%), and ALC-0159 (1.7%); all lipids and solvents were obtained from Merck. The aqueous phase consisted of mRNA diluted to 0.16 mg/ml in 50 mM citrate buffer (pH 4). Organic and aqueous streams were co-injected into the microfluidic mixer at a total flow rate of 10 ml/min with a 1:3 ethanol-to-aqueous flow-rate ratio, yielding nanoparticles with an N/P ratio of 6 and a lipid-to-mRNA mass ratio of 25:1. The resulting LNP suspensions were collected in RNase-free tubes and dialyzed using Slide-A-Lyzer™ dialysis cassettes (3.5 kDa MWCO; Thermo Scientific, Cat. 66330) against sucrose–phosphate buffer (150 mM sucrose, 75 mM NaCl, 10 mM sodium phosphate, pH 7.4), with buffer exchanges at 1 h, 2 h, and overnight. Final formulations were concentrated to 0.2 mg/ml using 100 kDa PES centrifugal filters (Sartorius Vivaspin, Cat. VS04T42), sterile-filtered through 0.22 µm PES membranes (Nalgene™, Cat. 720-1320), aliquoted, and stored at −80 °C until use.

### *Vaccine production: c*haracterization of mRNA–LNPs

mRNA–LNPs were characterized after a single freeze–thaw cycle. Encapsulation efficiency was measured using the RiboGreen assay (Invitrogen, Cat. R11491). LNP formulations (0.2 mg/ml mRNA) were diluted 1:50 in 1× TE buffer prepared from TE 20× stock (Invitrogen, Cat. T11493), and 50 µl was plated into wells containing either 50 µl of 1× TE buffer or Triton buffer (0.2% Triton X-100 prepared in 1× TE). After a 10 min incubation at 37°C, an equal volume (100 µl) of RiboGreen reagent diluted 1:100 in 1× TE was added, and fluorescence was recorded at 485/535 nm. Encapsulation efficiency (EE) was calculated using fluorescence values obtained from samples in 1× TE buffer (free mRNA, F_free) and Triton buffer (total mRNA, F_total) according to the equation EE (%) = [1 − (F_free / F_total)] × 100, and typically ranged from 89–96%. RNA content was interpolated from a standard curve (0.1–1.5 µg/ml) using the linear relationship F = A × C + B and corrected for the 200-fold dilution. Particle size and polydispersity index (PDI) were determined by dynamic light scattering using a Malvern Panalytical Zetasizer after diluting 50 µl of LNPs into 950 µl of storage buffer. Measurements were performed using a solvent refractive index of 1.338 and viscosity of 1.0547 mPa·s, with LNP optical parameters set to a refractive index of 1.46 and absorption of 0.01. Zeta potential was determined by electrophoretic light scattering after dilution of LNPs in 0.1× sucrose–phosphate buffer, using solvent parameters of refractive index 1.331, viscosity 0.8884 mPa·s, and dielectric constant 78.5.

#### Luciferase enzymatic assay

Luciferase assays were performed as previously described ^44^. For cell-based assays, cells were lysed using 1× Luciferase Cell Culture Lysis Reagent (Promega, Cat. E1531) prior to biochemical analysis. For tissue assays, one half of each organ was homogenized in lysis buffer and subjected to repeated freeze–thaw cycles, followed by centrifugation at 13,000 × g for 30 min to obtain the soluble protein fraction. Protein concentration was determined using the Bradford assay. Luciferase activity was measured in luciferase assay buffer, and relative luminescence units (RLU) were quantified in a 96-well plate using a Veritas luminometer (Turner Biosystems) with a 10-second integration time. Luminescence values were normalized to total protein content as determined by the Bradford assay.

#### Real-time PCR (RT-PCR)

Cells were lysed in TRIzol reagent (Thermo Fisher Scientific, Cat. 15596026), and total RNA was extracted from the aqueous phase using the Direct-zol RNA Miniprep Kit (Zymo Research, Cat. R2050) according to the manufacturer’s instructions. RT–PCR analyses were performed as previously described ^18^. Briefly, cDNA was synthesized using Moloney murine leukemia virus reverse transcriptase (Promega, Cat. M3681) and random primers (Promega, Cat. C118A). For each sample, negative control reactions were prepared in the absence of reverse transcriptase.

Quantitative RT–PCR was performed using SYBR Green chemistry (Promega, Cat. A600150), and cDNA samples were amplified in triplicate in 96-well plates using GoTaq qPCR Master Mix (Promega, Cat. A6001) in a final reaction volume of 10 μL, according to the manufacturer’s protocol, on a QuantStudio 3 96-Well 0.1 ml Block system (Thermo Fisher Scientific). The thermal cycling conditions were as follows: initial denaturation at 95°C for 2 min, followed by 40 cycles of 15 s at 95°C and 1 min at 60°C.

The primers used (Eurofins) were *Arg1* (5′-GAATCTGCATGGGCAACCT-3′; 5′-ACACGATGTCTTTGGCAGATAT-3′), *mChil3* (5′-TGGGCTAAGGACAGACCAAC-3′; 5′-ACTGAACGGGGCAGATCCAA-3′), and *Rplp0* (5′-GGCGACCTGGAAGTCCAACT-3′; 5′-CCATCAGCACCACAGCCTTC-3′). Gene expression levels were calculated using the comparative Ct method (2^−ΔΔCt^), with Rplp0 used as the housekeeping gene.

### Circulating cytokine levels measurement

Plasma collected from treated mice was stored at -80°C until use. At the time of the experiment, plasma samples were thawed, diluted two-fold in assay buffer and cytokine levels were measured by using the bead-based LEGENDplex Mouse Anti-Virus Response Panel (13-plex) immunoassay according to the manufacurer’s instructions (Biolegend, Cat #740622).

#### Generation and Treatment of BM-DCs

Bone marrow-derived dendritic cells (BM-DCs) were generated by culturing bone marrow cells for 8–10 days in dendritic cell (DC) medium consisting of RPMI (EuroClone, Cat. ECB9006L) supplemented with 10% fetal calf serum (FCS; Gibco, Cat. 10500–064), 10 ng/ml GM-CSF (PeproTech, Cat. AF-315-03), 100 U/ml penicillin (Orion, Cat. 465161) and 100 µg/ml streptomycin (Thermo Fisher Scientific, Cat.D7253), and 2 mM L-glutamine (Thermo Fisher Scientific, Cat. BP379–100). Cells were maintained at 37°C in a humidified atmosphere containing 5% CO_2_. Medium was supplemented or refreshed on days 3, 6, and 8. On day 8, cells were replated and treated on day 9 for 24 or 48 hours with 2.5 µg of mRNA formulated in LNPs or an equivalent amount of empty LNPs.

#### Flow Cytometric Analysis of BM-DC Maturation

To assess the maturation status of BM-DCs, the following fluorochrome-conjugated antibodies were used: CCR7-PE, MHC-II-APC-eFluor780, CD80-APC (BioLegend, Cat. 120105, clone 4B12, 47-5321-82, clone M5/114.15.2, 17-0801-81, clone 16-10A1) and CD86-FITC (BD Biosciences, cat. 11-0862-85, clone GL1). Fc receptor blocking (BD Pharmingen, Cat. 553142, clone 2.4G2) was applied in all experiments involving mouse cells. Unstained controls and fluorescence-minus-one (FMO) controls were included for each panel. Data acquisition was performed on a BD LSR Fortessa flow cytometer, and analysis was conducted using FlowJo software (Tree Star).

#### ELISA

The concentration of IL-12 in supernatants from BM-DCs was determined by ELISA using the Mouse IL-12/IL-23 p40 Allele-specific DuoSet ELISA kit (R&D Systems, Cat. DY499) following the manufacturer’s instructions. Briefly, 96-well MaxiSorp microplates (Thermo Fisher Scientific, Cat. 442404) were coated overnight with capture antibody. The following day, wells were washed and blocked with 1% BSA in PBS (Biowest, Cat. P6154; Lonza, Cat. 17-516F) for at least 1 h. After washing, standards and samples were added and incubated for 2 h. Detection antibody was then applied for 2 h, followed by a 20-min incubation with Streptavidin-HRP (R&D, Cat. DY998). Substrate solution (R&D, Cat. DY499) was added until visible colour developed (maximum 20 min), and the reaction was stopped with sulfuric acid (Acros Organics, Cat. 124645001, diluted 1:4 in water). Optical density was measured immediately at 450 nm, with 540 nm as the reference wavelength, using a Multiskan GO spectrophotometer (Thermo Fisher Scientific). IL-12 concentrations were calculated from the standard curve, and data analysis. All washing steps were conducted with three washes of 0.05% Tween-20 in PBS (Fisher BioReagents, Cat. BP337), followed by one wash with PBS alone.

Antigen-specific antibodies in sera from immunized mice were quantified by ELISA. Briefly, recombinant goat antigen, kindly provided by Prof. Ricagno, was diluted to a final concentration of 10 µg/ml in 0.05 M carbonate coating buffer, and 100 µl per well was used to coat 96-well plates overnight at 4 °C. Plates were blocked for 1 h at room temperature with PBS containing 1% (w/v) bovine serum albumin (BSA; Sigma-Aldrich, Cat. 1120180025). Serum samples from immunized mice were diluted 1:100 in PBS containing 1% BSA, added to the wells (100 µl), and incubated for 2 h at room temperature. Plates were washed four times with PBS containing 0.1% Tween-20. Horseradish peroxidase–conjugated anti-mouse IgG secondary antibodies (Bio-Rad, Cat. 1706516) were diluted 1:5,000 in PBS containing 0.1% BSA, added to the wells, and incubated for 1 h at room temperature, followed by four washes with PBS containing 0.1% Tween-20. Signal detection was performed using 3,3′,5,5′-tetramethylbenzidine (TMB; Thermo Scientific, Cat. 34028). The reaction was stopped with 2 N sulfuric acid, and absorbance was measured using a GloMax Discover microplate reader (Promega).

#### Statistics

Data are presented as the mean ± standard deviation unless otherwise noted in the figure legends. Statistical analyses were performed using Prism 7 (Version 8.00, GraphPad Software Inc.). Outliers were identified and excluded using the ROUT method with Q = 1. For comparisons between two groups, t-tests were applied. One-way ANOVA was used to evaluate differences among three or more independent groups, followed by Tukey’s post hoc test for pairwise comparisons or Dunnett’s test for comparisons against a control group. Two-way ANOVA, followed by Sidak’s post hoc test, was used to assess the effects of two factors in multiple comparisons. A p-value < 0.05 was considered statistically significant.

## Declarations

### Funding

The authors are grateful to the financial support from and PNRR M4C2-Investimento 1.4-CN00000041-PNRR_CN3RNA_SPOKE9 (to P.C.)., Research Council of Finland and Finnish Cancer Foundation (to S.F.)

### Author contributions

Conceptualization: E.B., N.R., A.V., C.S., S.P., C.D.V., L.O., D.M., S.F., S.M., E.V., P.C. ; methodology: E.B., A.P., N.R., C.M., C.S., S.P., F.A., M.L.G., C.M., G.C., M.R., C.D.V., L.V., L.O.; Investigation: E.B., A.P., N.R., C.M., C.S., S.P., F.A., M.L.G., C.M., G.C., M.R., C.D.V., L.V., L.O.; formal analysis: E.B., C.S., S.P., M.L.V., C.M., C.D.V., L.V., L.O.; Funding acquisition: P.C., S.F.; supervision: L.O., D.M., D.F., S.M., E.V., P.C..; writing of original draft: E.B., A.P., review and editing of the manuscript: L.O., D.M., D.F., S.M., E.V., P.C..; All authors read and approved the final manuscript.

### Competing interests

The authors declare no competing interests.

### Ethics Declarations

All animal experimentation was carried out in accordance with the Animal Research: Reporting of In Vivo Experiments (ARRIVE) guidelines and the European Guidelines for Animal Care. The animal study protocols were approved by: the Italian Ministry of Research and University (permission numbers: 417/2018, 416/2022, 904/2023, 105/2024), the Finnish Act on the Protection of Animals Used for Scientific or Educational Purposes (497/2013) and to the Directive 2010/63/EU of the European Parliament and of the Council of 22 September 2010 on the protection of animals used for scientific purposes and approved by the Finnish National Animal Experiment Board (Hankelupalautakunta – ELLA).

### Availability of data and materials

The data that support the findings of this study are available from the senior author upon reasonable request. The models are undergoing deposition at the Mutant Mouse Resource & Research Centers (MMRRC; www.mmrrc.org) NFkB-luc2, code:76124; NFkB-STOP, code: 76125; STAT6-STOP, code 76143; STAT6-luc2, code: 76144.

### Consent for publication

Not applicable.

### Disclosure statement

No potential conflict of interest was reported by the author(s).

## Notes

### Competing Interest Statement

The authors have declared no competing interest.

