## supplementary for "Spatiotemporal Atlas of Pro-Inflammatory (NF-κB) and Anti-Inflammatory (STAT6) Signalling Using Reporter Mice during mRNA Vaccination"

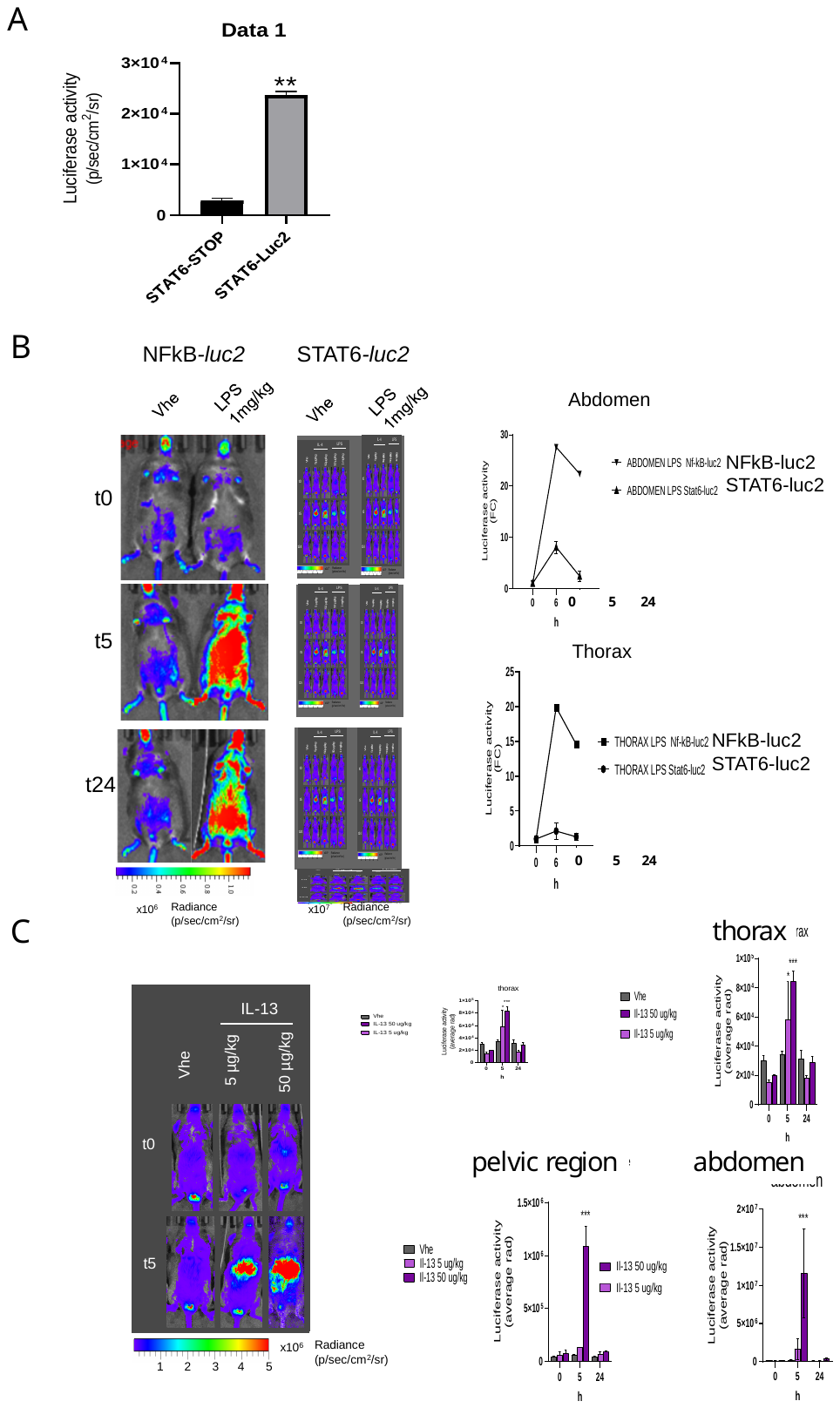

**Supplementary Figure 1 Validation of the STAT6 reporter mouse:**

**(A)** Quantification of whole-body bioluminescence (BLI) in STAT6*-*STOP and STAT6*-luc2* mice. Data are presented as total luciferase activity (n = 2). Statistical significance was assessed using an unpaired *t*-test (**p* < 0.01).

**(B)** *In vivo* imaging of NFκB*-luc2 and* STAT6*-luc2* mice treated with LPS (1 mg/kg, intraperitoneally) at 0, 5, and 24 hours post-injection. Representative images (left) show increased bioluminescence following LPS stimulation compared to vehicle-treated controls. Quantification (right) reports luciferase activity as fold change (FC) relative to baseline (t₀) for NFκB*-luc2* and STAT6*-luc2*. LPS administration induced a markedly stronger activation in NFκB*-luc2* mice compared with STAT6*-luc2* mice at equivalent doses.

**(C)** Validation of STAT6*-luc2* mice following intraperitoneal administration of 5 or 50 µg/kg of IL-13. BLI was quantified within defined regions of interest (Figure 2) and is reported alongside representative BLI images. Statistical analysis was performed using two-way ANOVA followed by Dunnett’s multiple comparison test versus vehicle (***p* < 0.001).

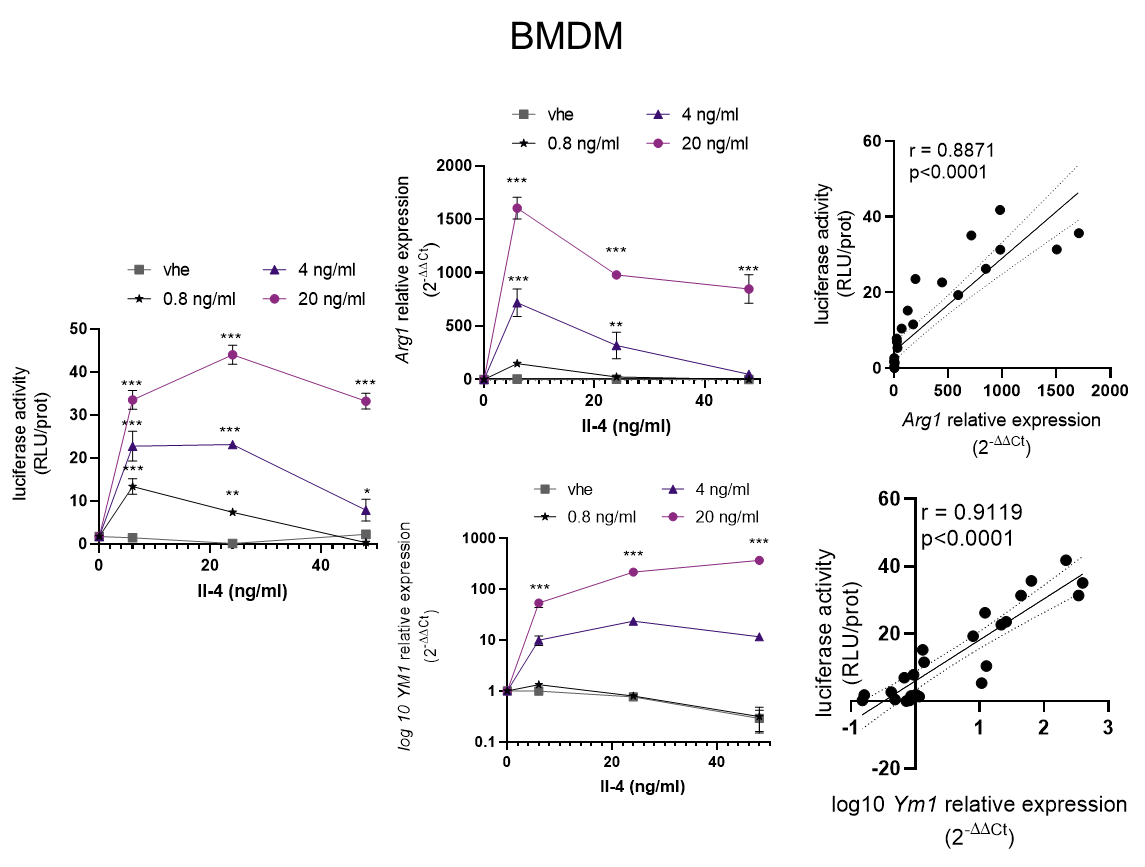

**Supplementary 2:** bone marrow derived macrophages (BMDM) were treated with IL-4 at different doses (0, 0.8, 4, 20 ng/mL) for 0, 6, 24, or 48 h. At each time point, luciferase expression in protein extracts was quantified and expressed as RLU per μg of protein ± SEM. At the same time points, relative quantification of target gene transcripts, *Arg1* and *Ym1* (*Chil3)* was performed using the 2^−ΔΔCt^ method versus vehicle (*Arg1* above, *Ym1* log10 expression below) ± SEM. Statistical analysis was performed using two-way ANOVA followed by Dunnett’s test versus vehicle (n = 2). Correlation between luciferase activity and gene relative expression is shown (right); Pearson r and p-values for each interpolated curve are reported.

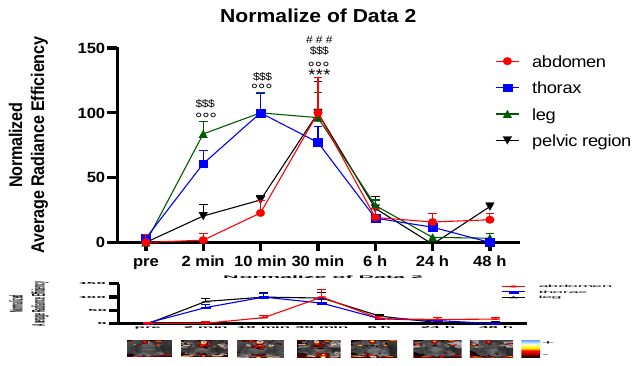

**Supplementary 3**: Kinetic profile of fluorescently labelled mRNA-LNPs *in vivo* in different body districts. C57BL/6 mice were intravenously injected with 10 µg of Cy5.5-labeled m¹Ψ-modified mRNA. Fluorescence was monitored at serial time points by *in vivo* imaging. Normalized fluorescence signals are presented as the mean ± SEM per mouse for each specified region of interest (ROI). For each body district, the highest detected signal has been placed at 100%. Representative *in vivo* images are shown.

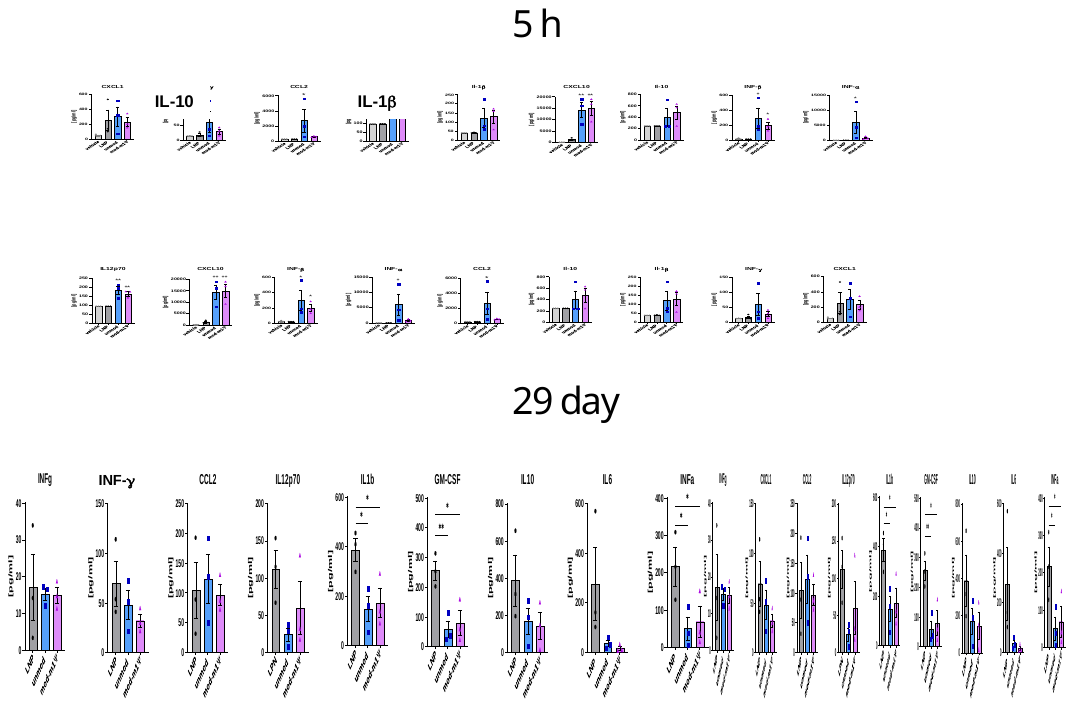

**Supplementary 4**: Cytokine levels in plasma were measured 5 hours after administration or at day 29 . Cytokine concentrations are presented as bar graphs ± SEM (n = 3).
